# Female-biased population sex ratios caused by genetic rather than ecological mechanisms

**DOI:** 10.1101/2023.12.07.569536

**Authors:** Xiaomeng Mao, Andres J. Cortés, Christian Rixen, Sophie Karrenberg

**Author notes:** **Corresponding authors:** Sophie Karrenberg.

## Abstract

1. Biased sex ratios among reproductive individuals are common in plants, but the underlying mechanisms, as well as the evolutionary consequences, are not well understood. The classical theory of Düsing and Fisher predicts an equal primary sex ratio at seed production, based on the selective advantage of the rare sex. Biased sex ratios among reproductive plants can arise from sexual dimorphism in survival and flowering. Sex ratio biases can also be present from seed; in these cases, assumptions of Düsing’s and Fisher’s theory, for example, random mating or demographic equilibrium, were not met.

2. We investigated mechanisms leading to female-biased sex ratios in the arctic-alpine dwarf willow *Salix herbacea* L. We studied sex ratios in three natural populations over three years as well as in 29 crosses (full-sib families) under controlled conditions over four growth periods. We tested whether sex ratio was associated with germination, survival or flowering, and whether females and males differed in habitat, size or flowering.

3. We detected a strong and consistent female bias, both in natural populations (sex ratio [proportion of females]: 0.71-0.82) and in our controlled experiment (overall sex ratio: 0.70-0-72). Our data did not support habitat segregation of the sexes or sexual dimorphism in size or flowering. Family sex ratios varied largely (from 0.25 to 1), including many female-biased families, but also unbiased families and two male-biased families. Families with lower germination, seedling establishment, survival or flowering did not have stronger female bias, indicating that intrinsically higher survival or flowering in females does not explain overall female bias.

4. Synthesis. Our results suggest that sex ratio bias in *S. herbacea* is already present in seeds and does not arise through intrinsic differences between sexes. Candidate mechanisms that can lead to both overall female bias and variation in sex ratio among families are meiotic drive or cyto-nuclear interactions. The pioneer habit of *Salix* may lead to non-equilibrium population dynamics that allow for the long-term persistence of variable genetic sex ratio distortion systems that arise from genetic conflict.

## Introduction

Sex ratio selection is one of the important forces implicated in the evolution of sex-determining systems, with effects on adaptation and lineage divergence, yet, mechanisms leading to sex ratio bias are generally not well understood (Barrett et al., 2010; Hardy, 2002; Uller et al., 2007). Darwin (1871), Düsing (1884; see Edwards, 2000 for English translation) and Fisher (1930) were already intrigued by sex ratios and proposed that females and males are expected to be equally common at birth or in seeds, due to a strong selective advantage of the rarer sex. Later theoretical treatments confirmed this theory and clarified assumptions under which the expectation of such equal primary sex ratios is valid; these include demographic equilibrium, random mating, biparental sex determination and Mendelian segregation of sex-determining variants (e.g., Bull, 1983; Bull & Charnov, 1988; Grafen, 2014; Hamilton, 1967; Hardy, 2002). Biases in the primary sex ratio have been attributed to mechanisms that violate these assumptions, for example, local mate competition rather than random mating, sex determination by only one parent (cytoplasmic sex determination), or unequal gamete transmission through either meiotic drive (non-Mendelian inheritance) or competitive advantage (Bull, 1983; Hardy, 2002; Meiklejohn & Tao, 2010; Unckless & Clark, 2014; Werren & Beukeboom, 1998). Even unbiased primary sex ratios often develop into biased sex ratios among reproductive individuals (secondary sex ratios), as a result of ecological differences between sexes (Barrett et al., 2010; Delph, 1999; Field et al., 2013). Overall, sex ratio biases evolve through both natural selection and genetic conflict, and the complex interactions between these processes are challenging to disentangle.

Separate sexes (dioecy) occur in about 6% of angiosperms with a wide phylogenetic distribution (Renner, 2014). Biased secondary sex ratios are common in dioecious plants, especially male bias, but the underlying mechanisms remain unresolved in most cases (reviewed in Barrett et al., 2010; Field et al., 2013; Sinclair et al., 2012). Sexual dimorphism in morphological and physiological traits can lead to sex-biased survival or flowering and thus biased sex ratios among reproductive plants (Barrett et al., 2010; Delph, 1999) as well as to spatial or habitat segregation of the sexes (Bierzychudek & Eckhart, 1988). For *Valeriana edulis*, for example, Petry et al. (2016) have shown that higher water use efficiency of females at low temperatures underlies increasing female bias at higher altitudes. Environmental determination of sex ratios may thus impact the distribution of dioecious species (Hultine et al., 2016; Petry et al., 2016). Sex ratio biases may also arise from seed (primary sex ratio bias). In *Rumex nivalis*, for instance, female determining pollen was more competitive than male-detemining pollen leading to more pronounced female bias in denser stands, consistent with local mate competition (Stehlik et al., 2008). However, one of the difficulties when studying plant sex ratios is that, unless molecular sex markers are available, sex can only be inferred morphologically once individuals are flowering, making it difficult to identify the stage at which sex ratio biases are generated (Barrett et al., 2010). Moreover, in clonal species, biases in the secondary sex ratio of ramets can result from higher ramet production of one sex (Field et al., 2013; Timerman & Barrett, 2019).

Willows (*Salix*) are one of the plant genera in which sex ratios have been frequently investigated. Almost all studies report female-biased secondary sex ratios in natural populations (Alliende & Harper, 1989; Bürli et al., 2022; Che-Castaldo et al., 2015; Crawford & Balfour, 1983, 1990; Dawson & Bliss, 1989; de Jong & van der Meijden, 2004; Dudley, 2006; Elmqvist et al., 1988; Hroneš et al., 2019; Hughes et al., 2010; Myers-Smith & Hik, 2012; Ueno et al., 2007). In part of these studies, sex ratios were associated with environmental conditions such as altitude, drought or the intensity of herbivory, and ecological mechanisms were implicated in the generation of secondary sex ratio bias (e.g., Bürli et al., 2022; Dawson & Bliss, 1989; Dudley, 2006; Elmqvist et al., 1988; Ueno et al., 2007). Studies on other *Salix* species, in contrast, did not find evidence for ecological differences between sexes and concluded that sex bias could be present from an early stage (Che-Castaldo et al., 2015; de Jong & van der Meijden, 2004; Hroneš et al., 2019; Myers-Smith & Hik, 2012). Data on the seed sex ratio, however, is only available for a few *Salix* species, including *S. repens* and *S. viminalis*, where an overall female bias in seed families was found suggesting that the bias might arise from an early stage in these species (Alström-Rapaport et al., 1997; de Jong & van der Meijden, 2004).

Here we contribute a study on the mechanisms leading to female-biased sex ratios in the arctic-alpine dwarf willow *Salix herbacea* L. (2-10cm). For this species, female bias among reproductive individuals has been described for populations in Iceland (Crawford & Balfour, 1983) and Switzerland (Wheeler et al., 2016). *Salix herbacea* is long-lived and reproduces both clonally and sexually (Beerling, 1998; Centenaro et al., 2023). The species forms intermingled mats with underground stolons, such that individuals cannot be identified in the wild (Beerling, 1998; Centenaro et al., 2023).

We investigated sex ratios in three natural populations of *S. herbacea* over three years as well as in crosses (full-sib families) that we cultivated under controlled conditions for four growth periods. We tested whether our data is consistent with four hypotheses on the generation of female bias among reproductive plants, assuming unbiased seeds (Table 1): **(1)** higher germination, **(2)** higher survival, or **(3)** more flowering in females as compared to males, as well as **(4)** more apparent females, that are more likely to be sampled. To this end, we tested whether the two sexes differed in the main habitat parameters in their environment, snowmelt time and elevation, which would indicate differential survival between sexes. In our controlled experiment, we tested for environmentally independent associations of family sex ratios with germination, survival and flowering. In addition, we analyzed whether females were larger or flowered earlier than males, and thus were more apparent. Using data from both a field survey and a controlled experiment allows us to disentangle possible mechanisms for the generation of female bias.

**Table 1.**
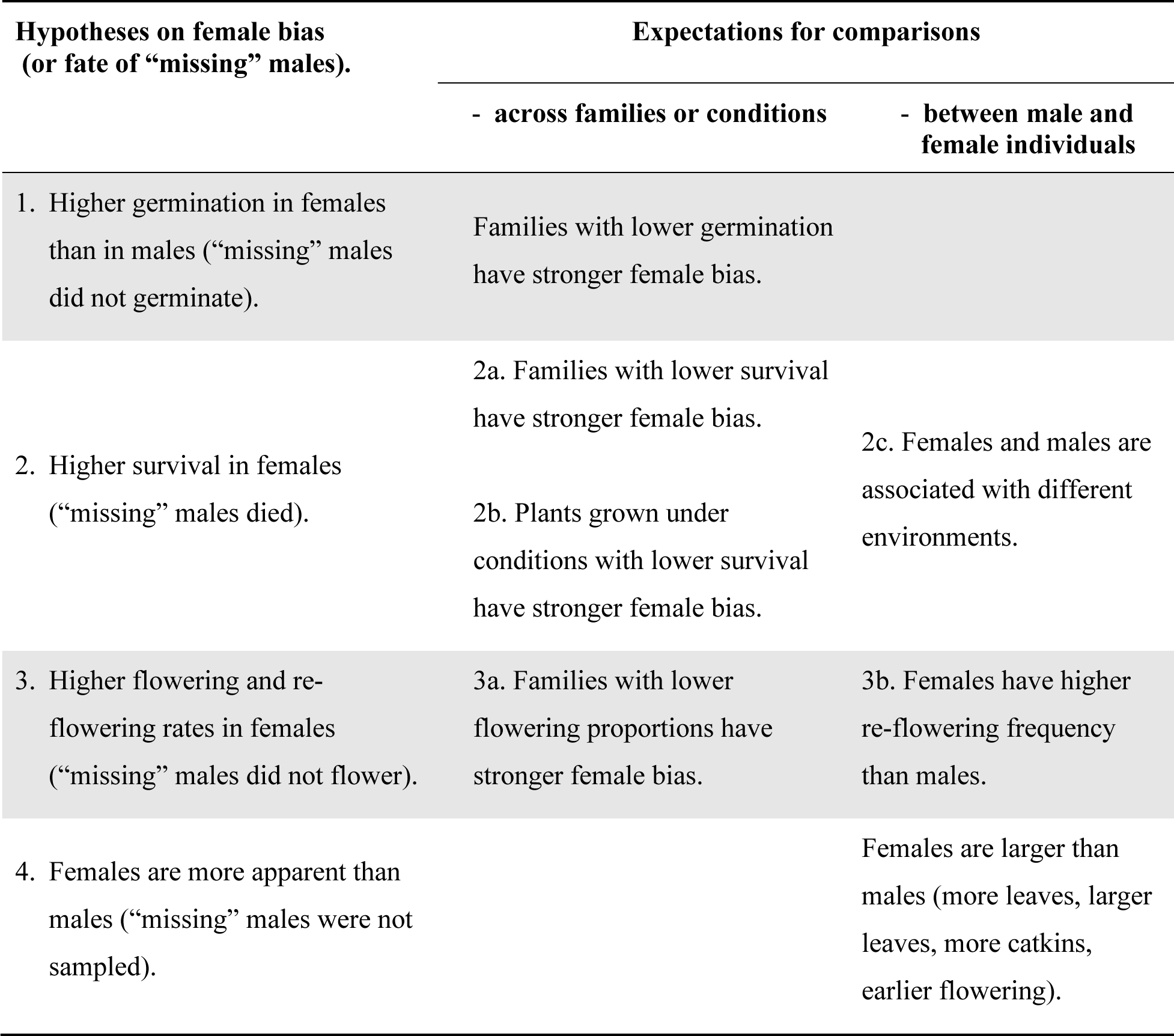
Four hypotheses on the generation of female bias after seed production, assuming equal sex ratio at seed production.

## Materials and Methods

### Natural populations: sex ratio and differences between sexes in habitat and morphology

Dwarf willow occurs above the treeline in the Alps and at high latitudes (arctic-alpine distribution). In this environment, the growing season is only 3-4 months long; for the rest of the year, plants are exposed to freezing temperatures or covered by snow (Beerling, 1998; Wheeler et al., 2016). We re-analyzed published data from three natural *S. herbacea* populations near Davos Switzerland, at elevations from 2000 to 2800 m above sea level (Wheeler et al., 2016), supplemented by unpublished records from the same study (see below). A total of 120-124 round patches of *S. herbacea* with 10 cm diameter per population (484 patches in total) were marked along altitudinal transects before flowering (Wheeler et al., 2016). Plant sex in each patch (as female, male or both sexes) was recorded over three years (2011-2013), at approximately weekly visits to each site during the flowering period (May-July), and the start of flowering was noted as days after snowmelt (Wheeler et al., 2016). In the second year, we estimated a size proxy, leaf area, as leaf width multiplied by leaf length, averaged for two fully expanded leaves on flowering twigs. We further recorded two of the main environmental gradients in this environment for each patch: elevation and snowmelt time (as day of year, in each year, Wheeler et al., 2016).

We estimated the sex ratio as the proportion of females among reproductive plants in each year using 349 patches where only one sex was observed, as well as 135 patches that exhibited flowering ramets of both sexes, either in the same year or across years, and were excluded from Wheeler et al. (2016). Patches with flowering individuals of both sexes in the same year were included in the calculation of sex ratio as 0.5 individuals for each sex that year. We tested whether the sex ratio in each year and population deviated from 0.5 using a *z*-test (*prop.test* command) in R. All analyses were conducted in R version 4.3.1 (R Core Team, 2022).

We used linear mixed models to test whether elevation and mean snowmelt time across years differ between patches with male, female or both sexes flowering, and whether flowering time and leaf area, both measured in the second year, differ between males and females. In these models, patch sex was a fixed effect and population was a random effect. Linear mixed models were performed using *lmer* of the R package *lme4* (Bates et al., 2015) and significance tests using *lmerTest* (Kuznetsova et al., 2017).

### Controlled conditions: sex ratio and difference between sexes in full-sib families

We generated 29 hand-pollinated crosses in a natural alpine population of *S. herbacea* near Jakobshorn, Switzerland (46.7720N, 9.8554E, 2535 m above sea level) between June and August 2020. Pre-receptive catkins on female individuals were bagged with dialysis tubing (Karrenberg & Suter, 2003) to exclude natural wind and insect pollination. Each female was later pollinated using male catkins from a single individual, collected on the same day. Each male was used for one cross only. Mature fruits were collected 4-6 weeks after pollination. Individuals used for crosses were at least several meters apart.

A total of 1521 seeds from 29 families (seed number per family: 52.45 ± 9.07 [mean with standard error], see also Supplementary Table 1) were placed on Petri dishes with wet filter paper and kept at 0° C without light for three months to break seed dormancy (unpublished results of the authors). We transferred seeds to a growth chamber at 20 °C with 16 hours of light for germination (7-30 days). A total of 1428 seeds germinated, and were planted into 7×7×8 cm pots filled with a 1:1 (by volume) mix of Rhododendron soil (Plantasjen, Kongsvinger, Norway) and clay balls (Leca Sverige AB, Linköping, Sweden), covered by a top layer of gravel. 1288 individuals were arranged in a randomized block design (8 blocks with 6×9 pots on each of three tables), comprising 46 ± 8.05 individuals per family (Supplementary Table 1), in a greenhouse at Uppsala University (20°C, day and night, 16h, with supplementary light). Established seedlings (26-46 days after planting) were moved to a climate-controlled chamber in a compacted randomized block design, where empty spots of the first two tables were filled with survivors of the third table. Further 55 empty spots were filled with additional established seedlings from the same experiment, yielding 757 established seedlings in total (26.10 ± 4.20 per family, Supplementary Table 1). We watered plants as needed and applied all-purpose liquid fertilizer 3-4 times per growth period (Formulex, Growth Technology, Taunton, United Kingdom).

We cultivated plants for four growth periods (2020-2023), with artificial spring/summer (35–49 days, 14–16°C / 4-10°C day/night, 14–16 h light) and autumn (70– 80 days, temperatures and light decreasing from 16°C / 10°C day/night and 16h light to 6°C /4 °C day/night and 12h of light, Supplementary Figure 1). The temperature and light scheme were designed to be similar to maximum temperatures and day length in the source population (Supplementary Figure 1). After bud set and leaf shedding under autumn conditions, we exposed plants to a shortened, artificial winter at 0°C without light (as under snow cover) for 120-180 days, such that one entire growth cycle could be completed within 7-8 months (Supplementary Figure 1).

In each growth period, we checked plants for flowering regularly and assessed plant sex. In growth period four, we recorded bud break, leaf expansion and flowering every three days, counted the number of leaves and catkins and measured the length and width of two leaves per plant, as well as the length of the longest twig.

We calculated the proportion of reproductive individuals and the sex ratio (proportion of females) among reproductive individuals, in each growth period and overall, i.e., including all individuals that flowered at least once. We tested whether overall family sex ratios deviated from equal sex ratios using a *z*-test (*prop.test* command) in R, as described above. We analyzed whether the family sex ratios were affected by proportions of germination, seedling establishment, adult survival or flowering (hypotheses 1-3, Table 1) using linear regression analyses weighed by the number of seeds, seedlings, or adult plants per family. For these analyses, we excluded three families with fewer than three surviving individuals (Supplementary Table 1). We used mixed models with binomial errors (*glmer*) to test whether females and males differed in the probability of re-flowering (hypothesis 3) using the packages *lme4* and *lmerTest* in R (Bates et al., 2015; Kuznetsova et al., 2017). We used mixed models with gaussian errors (*lmer*) to test whether females and males differed in phenological timing (bud break, leaf expansion, and flowering) or morphological traits (leaf area [log-transformed], leaf number, length of the longest twig, and catkin number, hypothesis 4) using the same R packages. In these models, plant sex was a fixed effect, and block and family were random effects.

We used 35 females and 28 males to generate crosses for additional experiments not included here, and these individuals were temporarily placed in a different growth chamber with similar conditions. These individuals did not affect the results of our statistical analyses.

## Results

### Sex ratio in natural populations and under controlled conditions

Natural populations exhibited strong female bias: the sex ratio (proportion of females) among reproductive plants varied between 0.71 and 0.82 in the three populations and three years of observation (Figure 1A). In all cases, the sex ratio deviated significantly from 0.5 (Figure 1A).

**Figure 1.**
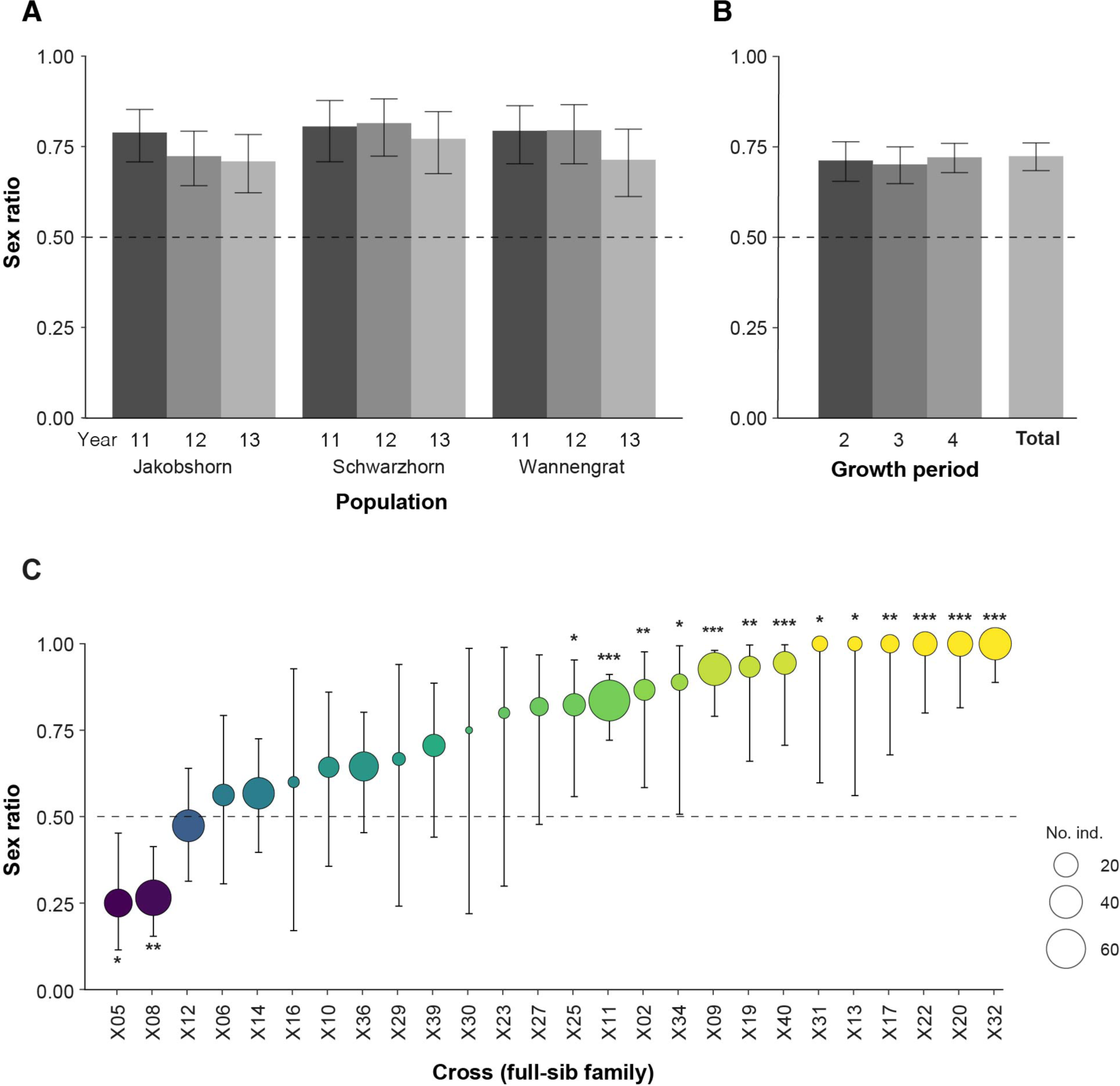
Sex ratios (proportion females) among reproductive individuals of the dwarf willow *Salix herbacea* with 95% confidence intervals in three natural populations in Switzerland (J: Jakobshorn, S: Schwarzhorn, W: Wannengrat), 2011-2013 (**A**), in a controlled experiment over three growth periods, as well as overall (**B**) and in 26 families of the same experiment (**C**). Sex ratios in A and B were all significantly higher than 0.5 (z-test). In C, the results of z-tests against 0.5 are indicated as *** P < 0.001, ** P < 0.01 and * P<0.05. The dashed line indicates an equal sex ratio of 0.5.

In our experiment, 1428 of the 1521 seeds germinated (overall germination proportion 0.94) and were planted. Of these, 757 seedlings survived the establishment phase (overall proportion of seedling establishment: 0.53), and 728 individuals survived to the end of the experiment after four growth periods (overall proportion of adult survival: 0.96, Supplementary Table 1).

Plants started flowering from growth period 2 (flowering proportion: 0.37). The flowering proportion increased over subsequent growth periods (0.44 and 0.67 in growth periods 3 and 4, Supplementary Table 2). In total, 550 individuals flowered at least once, yielding an overall flowering proportion of 0.76. All individuals that flowered multiple times (199 individuals flowered twice, 172 individuals flowered three times) had the same sex across years. One individual exhibited deformed sexual organs that combined male and female traits in the same flower and was excluded from further analyses. Of the 28 plants that died after seedling establishment, 27 had not flowered and one was female (Supplementary Table 1). Another female of family X11 was removed from the experiment for tissue sampling.

The sex ratio of flowering plants (proportion of females) was stable across growth periods (0.71, 0.70, and 0.72 in growth periods 2, 3, and 4, respectively) with an overall sex ratio of 0.72 (Figure 1B, Supplementary Table 2), similar to the sex ratio in natural populations (Figure 1A). Sex ratios within families, however, varied strongly from 0.25 to 1. Half of the 26 families had sex ratios significantly higher than 0.5, including 6 families with females only (Figure 1C, Supplementary Table 1). Surprisingly, two families (X05 and X08) presented significant and strong male bias with sex ratios of only 0.25 and 0.27. For the remaining families, we did not find evidence for significant deviations in the sex ratio from 0.5.

### Tests for processes underlying sex ratio bias

#### Germination (hypothesis 1)

In our experiment under controlled conditions, the germination proportion within families ranged from 0.83 to 1 (Figure 2A). Female proportion increased slightly, but significantly, with germination proportion (Figure 2A, Table 2). However, this result does not support variation in germination underlying female bias, where a decrease in female proportion would be expected at higher germination rates.

**Figure 2.**
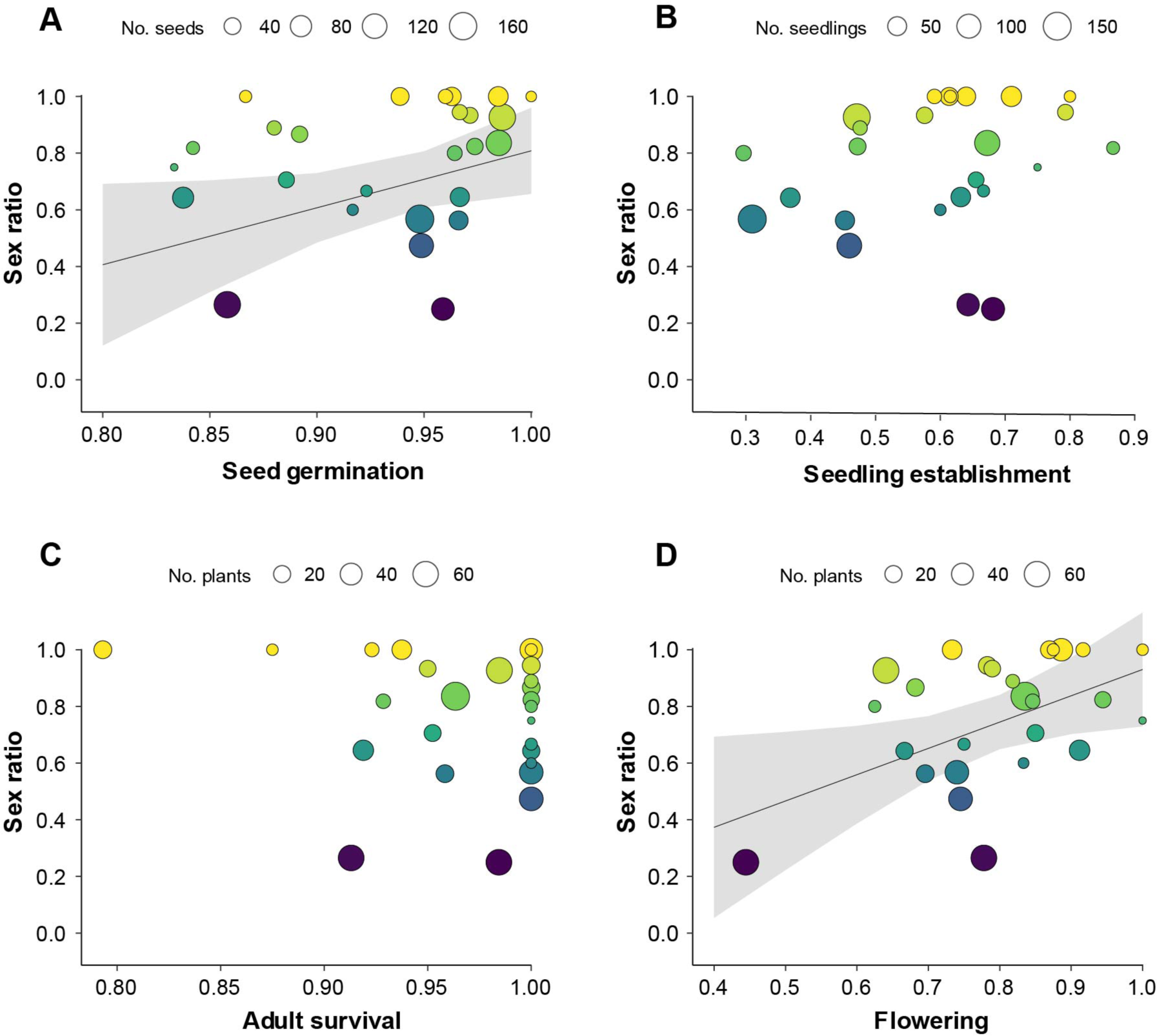
Relationship between family sex ratio (proportion females) and proportions of seed germination (**A**), seedling establishment (**B**), adult survival (**C**), and overall flowering (**D**) in the dwarf willow *Salix herbacea*, grown under controlled conditions. Linear regressions weighed by family size were significantly A and D, where regression lines with 95% confidence intervals are given.

**Table 2.** Linear regression results for sex ratio (proportion of females) in families of the dwarf willow *Salix herbacea*, as affected by the germination, seedling establishment, survival, and flowering. Regression analyses were weighed by family size. * P-Value < 0.05.

| Predictor | F-ratio | P-Value |
| --- | --- | --- |
| Seed germination | 4.63 | 0.04 * |
| Seedling establishment | 0.49 | 0.49 |
| Survival | 0.03 | 0.86 |
| Flowering | 5.66 | 0.03 * |

#### Survival and habitat association (hypothesis 2)

Seedling establishment per family ranged from 0.30 to 0.87 (overall 0.53) and was not associated with sex ratio (Figure 2B, Table 2), speaking against the involvement of differential survival in generating female bias (hypothesis 2a). Survival of adult plants was generally high (over 0.79 per family) and no association was found with sex ratio (Figure 2C, Table 2).

We observed large and significant differences in seedling establishment across the three main original blocks (tables) of our randomized block design, with proportions of established seedlings of 0.60, 0.60 and 0.44 on tables 1, 2 and 3 (*X^2^* = 30.96, df = 2, p-value = 1.90 × 10^−7^). The sex ratio, however, did not differ significantly between tables (0.72, 0.74 and 0.70 on tables 1, 2, and 3; *X^2^* = 0.58, df = 2, p-value = 0.75). These findings fail to support hypothesis 2b where a decreased sex ratio is expected on the table with the highest survival.

Our data from natural populations provides little evidence for habitat differences between sexes (hypothesis 2c): patches with males, females, or both sexes were found at elevations ranging from 2109m to 2780m (Figure 3B). Patches with females had slightly higher mean elevations (2455 m a.s.l.) than patches with males or both sexes (2412 and 2412 m a.s.l., Fig. 3B); this difference was marginally significant (Table 3). Snowmelt time ranged from day of the year 120 to 205 (means near 158, Figure 3A) and no significant difference between patches with females, males or both sexes was detected (Table 3).

**Figure 3.**
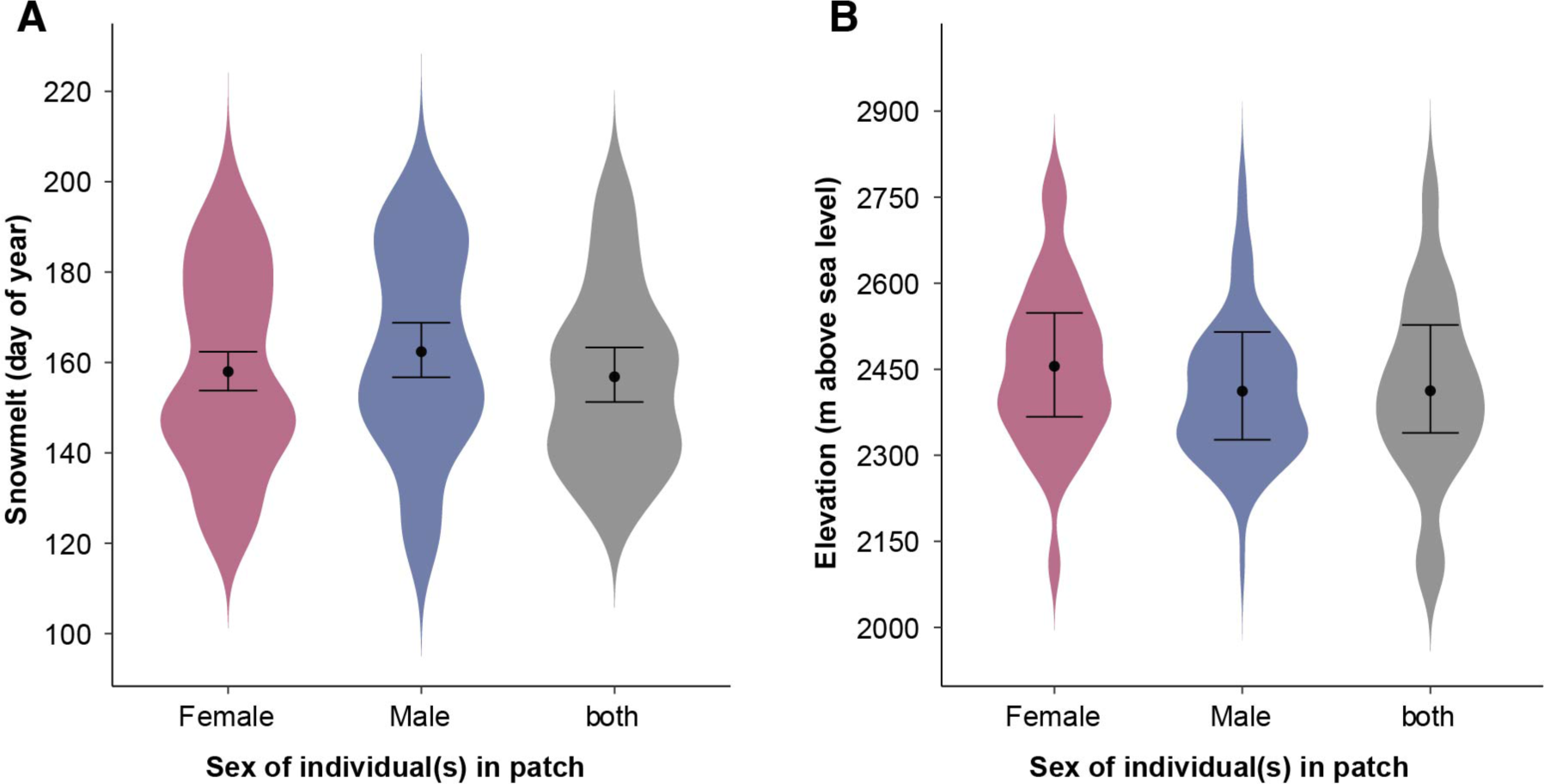
Snowmelt days and elevation in *Salix herbacea* patches in natural populations in Switzerland with female, male or both sexes flowering. Means and 95% confidence intervals are indicated as points and T-bars.

**Table 3.** Mixed model analyses of elevation, snowmelt, leaf area and day of flowering after snowmelt, as affected by the sex of dwarf willow (*Salix herbacea*) in three populations near Davos, Switzerland. Plant sex was a fixed factor and population was a random factor. Elevation and snowmelt were analyzed for 10 cm patches with male individuals, female individuals or individuals of both sexes. Leaf area and flowering day were measured on flowering twigs within patches and were analyzed for male and female individuals.

|  | **F** | **nominator df** | **denominator df** | **P-Value** |
| --- | --- | --- | --- | --- |
| Elevation | 2.94 | 2 | 407.18 | 0.05 |
| Snowmelt day<sup>a</sup> | 1.59 | 2 | 429.8 | 0.2 |
| Leaf area (log) | 0.75 | 1 | 304.56 | 0.39 |
| Day of flowering | 0.92 | 1 | 276.44 | 0.34 |

<sup>a</sup> averaged over three years

#### Flowering (hypothesis 3)

Families differed strongly in the proportion of reproductive individuals, ranging from 0.44 to 1 (Figure 2D, Supplementary Table 1). The sex ratio increased significantly with the proportion of reproductive individuals (Figure 2D, Table 2). If sex biases in flowering contributed to female bias, we would expect a contrasting pattern, an increase in sex ratio with decreasing flowering proportions, where more males (“missing males”) would remain vegetative (hypothesis 3a, Figure 2D).

Males and females re-flowered similarly: between growth periods 2 and 3, the re-flowering proportion was 0.63 in females and 0.66 in males; between growth periods 3 and 4, the reflowering proportion was 0.91 in females and 0.95 in males. Our mixed model analyses did not support differences between females and males in re-flowering (Table 4). We therefore find no evidence of more frequent flowering in females as a potential mechanism leading to female bias (hypothesis 3b).

**Table 4.** Mixed model analysis of survival, re-flowering probability, leaf area, leaf number, length of the longest twig, catkin number and day of flowering as affected by the sex of dwarf willow (*Salix herbacea*) grown under controlled conditions. Plant sex was a fixed effect, family and experimental blocks were random effects. Re-flowering probability was analyzed using models with binomial errors and a ꭓ^2^-test (model comparisons), whereas the remaining traits were analyzed using models with gaussian errors and F-tests with Satterthwaite’s correction. nom. df, nominator df; denom. df, denominator df. ** P-Value < 0.01, *** P-Value < 0.001.

|  | **X<sup>2</sup> / F-ratio** | **df /**<br>**nom. df, denom. df** | **P-Value** |
| --- | --- | --- | --- |
| Re-flowering probability<br>period 2 to 3 | 0.66 | 1 | 0.42 |
| Re-flowering probability<br>period 3 to 4 | 1.35 | 1 | 0.25 |
| Leaf area (log) | 1.06 | 1, 517.58 | 0.3 |
| Leaf number | 0.63 | 1, 540.83 | 0.43 |
| Length of the longest twig | 3.78 | 1, 522.19 | 0.05 . |
| Day of bud break | 1.62 | 1, 546.10 | 0.20 |
| Day of leaf expansion | 9.56 | 1, 523.60 | 2.10×10<sup>-3</sup> ** |
| Day of flowering | 13.19 | 1, 486.44 | 3.11×10<sup>-4</sup> *** |
| Catkin number | 32.73 | 1, 531.11 | 1.77×10<sup>-8</sup> *** |

#### Observation bias (hypothesis 4)

The mean leaf area of reproductive individuals did not differ between sexes: leaf area was 67.82 ± 0.08 and 71.60 ± 0.10 mm^2^ in natural populations and 165.67 ± 0.03 and 171.06 ± 0.04 mm^2^ in our experiment, for female and male individuals, respectively (mean with standard error, Figure 4F, Table 3 and 4). We found no evidence for differences between sexes in leaf number either (93.21 ± 4.35 and 95.78 ± 4.92 for females and males, hypothesis 4, Figure 4D, Table 4). The length of the longest twig was 35.06 ± 1.37 mm for females and 32.61 ± 1.64 mm for males with marginally significant differences between sexes (Figure 4G, Table 4).

**Figure 4.**
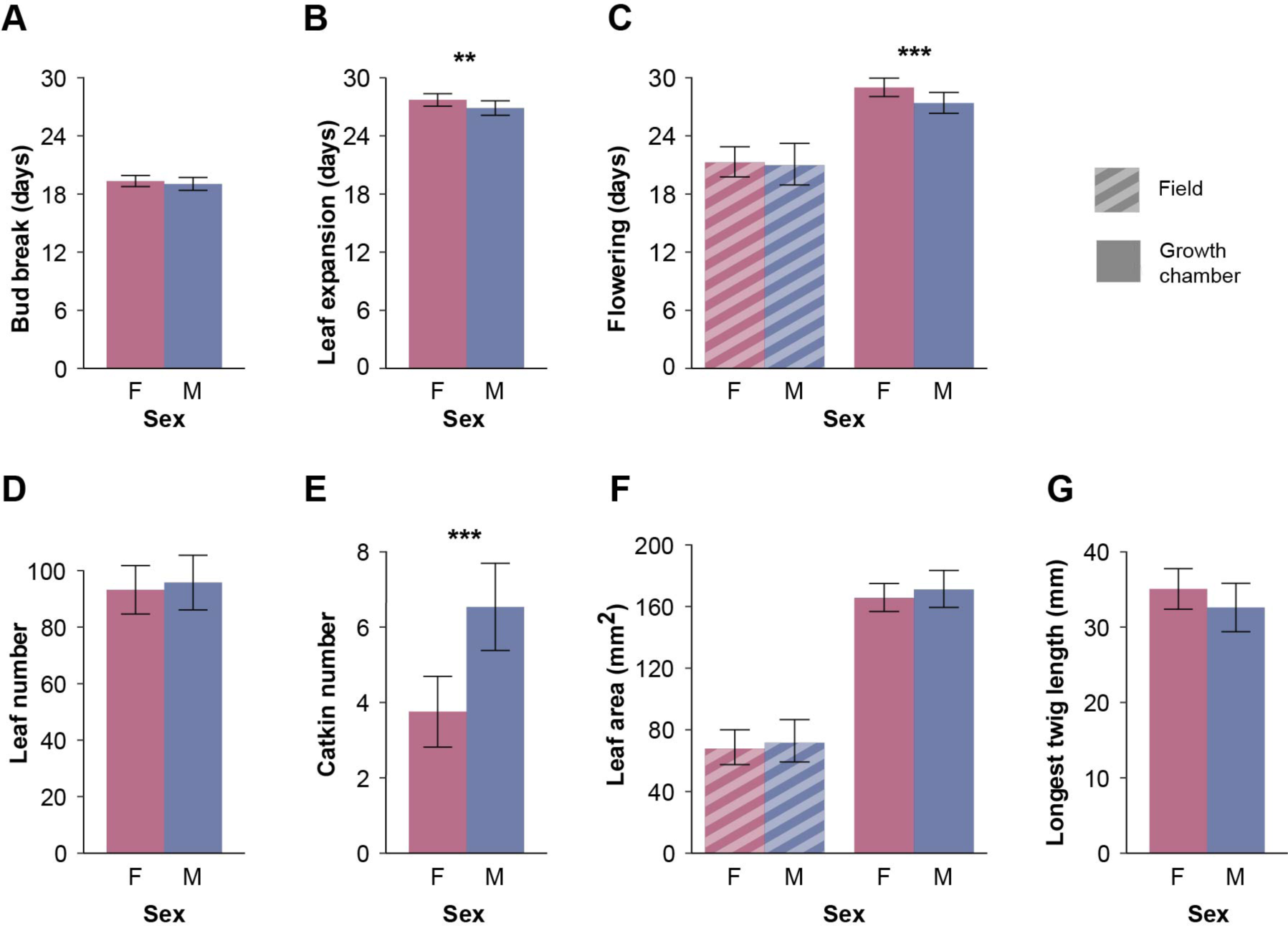
Sexual dimorphism in the dwarf willow *Salix herbacea* in natural populations in Switzerland (hashed bars) and in an experiment under controlled conditions (full bars): days of the year after snowmelt (natural populations) or days after the onset of spring conditions (experiment) for bud break (**A**), leaf expansion (**B**), and flowering (**C**), and length of the longest twig (**D**), catkin number (**E**), leaf number (**F**), and leaf area (mm^2^) (**G**). Significant differences between females (F) and males (M) are indicated as *** P< 0.001, **P<0.01.

Females and males started flowering 24.52 and 23.09 days after snowmelt in natural populations, again without evidence for a difference between sexes (Figure 4C, Table 3). Different from the field study, females flowered later than males in our experiment (on average, 29.34 and 27.61 days after the start of the spring/summer conditions for females and males, respectively, Figure 4C, Table 4). Females also had later leaf expansion time than males (27.72 and 26.87 days, Figure 4B, Table 4), but similar bud break time as males (19.34 and 19.04 days, Figure 4A, Table 4). Moreover, the catkin number in our experiment was 3.75 in females, significantly lower as compared to males (6.54, Figure 4E, Table 4). Our data thus do not support the hypothesis that more apparent females could contribute to estimates of female bias in this study.

## Discussion

### Sex ratios are strongly female-biased in natural and experimental populations

The sex ratio of flowering plants of *Salix herbacea* was strongly female-biased, with a proportion of females of 0.71-0.82 in three natural populations in its alpine environment above the treeline. Strong female bias (0.65-0.80) has also been observed in *S. herbacea* populations in Iceland (Crawford & Balfour, 1983) and in natural populations of many other *Salix* species in various habitats, including dwarf or small shrubs in arctic or alpine habitats (Crawford & Balfour, 1990; Dawson & Bliss, 1989; Myers-Smith & Hik, 2012) and small to medium shrubs in boreal or montane/sub-alpine habitats (Bürli et al., 2022; Che-Castaldo et al., 2015; Dudley, 2006; Elmqvist et al., 1988; Hroneš et al., 2019) as well as in temperate habitats (Alliende & Harper, 1989; de Jong & van der Meijden, 2004; Ueno et al., 2007). In our study, we further cultivated crosses of *S. herbacea* under controlled conditions, at temperatures near maximum daily temperatures in the field (4-16 °C, 3-4 months growth period). *Salix herbacea* grew well under these artificial conditions and produced about twice as large leaves, on average, as compared to plants growing in natural environments. Under these benign conditions, the overall sex ratio among flowering plants was also female-biased, 0.70-0.72, similar to the sex ratio in natural populations. The sex ratio was also constant over the three main blocks of the experiment, despite strong variation in seedling establishment across blocks (hypothesis 2b, Table 1). Sex ratios of plants grown from seeds are only rarely reported together with population sex ratios. In *S. repens*, open-pollinated seed families also exhibited female-biased sex ratios, as in wild populations (de Jong & van der Meijden, 2004). A consistent sex ratio bias under both benign and stressful conditions, as observed here in *S. herbacea*, strongly speaks against environmentally mediated survival differences between sexes as the main cause of the bias. Rather, sex ratio bias must arise either through intrinsic differences between sexes in germination, mortality, or flowering, to be discussed in the next sections, or be present from seed.

### Males and females are ecologically and morphologically similar

Our interpretation that the female-biased sex ratio of *S. herbacea* does not result from ecological differences between the sexes is further strengthened by direct observations of ecological and morphologic similarity in females and males. The natural habitat of *S. herbacea* is highly heterogeneous; differences in snowmelt timing and elevation lead to growth period variation from as little as 3 months to up to 5 months (Wheeler et al., 2016). Patches with males and females did not differ in snowmelt timing, but there was a (marginally significant) tendency for higher elevation in patches with females. Our data thus provides little support for spatial or environmental segregation between sexes (hypothesis 2c). Neither did we find evidence for secondary sexual dimorphism: females and males did not differ in leaf number or leaf size in our experiment. Previous studies in natural populations of *S. herbacea* did not detect differences between females and males either, in terms of phenology, leaf damage by herbivores and fungi, and carbon balance (Sedlacek et al., 2015; Wheeler et al., 2016). Similar size in both sexes further makes it unlikely that oversampling of females contributed to our observation of female bias (hypothesis 4). However, females had slightly longer twigs than males, most likely related to bearing fewer but heavier catkins with fruits than males. A lack of spatial or habitat segregation and sexual dimorphism was reported for a number of other *Salix* species (Alliende & Harper, 1989; Che-Castaldo et al., 2015; Hroneš et al., 2019; Ueno et al., 2007). In further *Salix* species, however, female bias was also present in almost all populations but varied along biotic or abiotic environmental gradients (Bürli et al., 2022; Dawson & Bliss, 1989; Dudley, 2006; Elmqvist et al., 1988). In many female-biased *Salix* species, sexes differ in stress tolerance (e.g., Dawson & Bliss, 1989; Xia et al., 2023) and pathogen damage (e.g., Alliende, 1989; Boecklen et al., 1990; Moritz et al., 2016). It is unclear though, whether these differences contribute to female bias or, alternatively, are a consequence of physiological differences between sexes. In *S. herbacea* there may also be slight differences in sex ratios along environmental gradients, such as elevation, but our results suggest that environmental factors are not the main cause of sex ratio bias.

### Germination, survival or flowering are not associated with sex ratio

Females and males were ecologically and morphologically similar; however, female bias could be the result of intrinsically higher survival or flowering in females (hypotheses 1-3). Our results do not support any of these hypotheses. Families with lower germination, seedling establishment, adult survival or flowering did not have stronger female bias, as would be expected if the “missing males” were among the individuals that died or did not flower. In our study, germination proportion and adult survival were high (0.79-1), whereas, at the seedling stage, nearly half of the individuals died. This indicates that there was ample opportunity for intrinsic differences between males and females to be detected, even under benign experimental conditions. The sex ratio in our experiment was stable over the three growth periods and the re-flowering probability did not differ between sexes. This suggests that males and females reached sexual maturity at a similar early age (2-3 years) in *S. herbacea*. Che-Castaldo et al. (2015) also found no evidence of biased mortality or flowering in *S. sitchensis*. Ueno et al. (2007) found no sex difference in mortality either, but in their study on *S. sachalinensis,* sex ratios were unbiased in young populations and female-biased in mature populations, suggesting that females reach maturity later than males. In our study, we find no evidence that sex ratio bias arises through biased germination, survival or flowering in *S. herbacea*, and this suggests that the sex ratio is already female-biased in seeds.

### Mechanisms potentially generating sex ratio bias

Even though the overall sex ratio had a strong female bias, the different families varied largely in sex ratio. Interestingly, both de Jong and van der Mejden (2004) and Alström-Rapaport et al. (1997) also found highly variable sex ratios in seed families of *S. repens* and *S. viminalis*, respectively. Mechanisms that can generate female bias include, for example, gametic competition, sex chromosome meiotic drive, and cyto-nuclear conflict (Barrett et al., 2010; Hurst & Werren, 2001; Jaenike, 2001; Meiklejohn & Tao, 2010; Werren & Beukeboom, 1998). Primary sex ratio bias through pollen competition, as detected in *Rumex nivalis* (Stehlik et al., 2008) requires a male heterogametic sex determination system where pollen is male- or female-determining. *Salix herbacea*, however, has a ZW sex determination system, where ovules produced by females are sex determining (unpublished results from the authors), as is common in *Salix* species from the subgenus *Vetrix*, such as *S. pupurea* and *S. viminalis* (Pucholt et al., 2015; Zhou et al., 2018). We can therefore exclude gametic competition as a mechanism generating sex ratio bias in *S. herbacea*. Both female meiotic drive and cyto-nuclear interactions are expected to elicit the evolution of sex ratio restorers that segregate in populations and lead to variation in sex ratio among families (Meiklejohn & Tao, 2010; Werren & Beukeboom, 1998), as we have detected here. Meiotic drive and cyto-nuclear interactions are thus candidate mechanisms underlying female bias in *S. herbacea* and potentially other *Salix* species.

Sex ratio theory predicts equal sex ratios and the main argument is that populations with biased sex ratios can be invaded by variants that lead to higher production of the rare sex until an unbiased sex ratio is reached (Düsing, 1884; Edwards, 2000; Fisher, 1930). It is an exciting speculation whether this mechanism gave rise to the male-biased families in *S. herbacea* in our study, that exist in a population with a strong female bias. Interestingly, pollen limitation of seed set was reported for *S. herbacea* (Finderup Nielsen, 2014), as well as in other *Salix* species (Fox, 1992; Totland & Sottocornola, 2001). Pollen limitation of seed set may result from female sex ratio bias and further increases the selective advantage of producing male offspring. In general, one of the situations where biased sex ratios can prevail, despite sex ratio selection, are non-equilibrium populations that have not reached a stable age distribution (Grafen, 2014). *Salix* species are pioneers that colonize new habitats with wind-dispersed seeds, for example, flood plains and glacier forefields (Karrenberg et al., 2002) such that a stable age distribution may never be reached. In *S. herbacea*, individuals also reproduce clonally and can be very long-lived, potentially thousands of years (Centenaro et al., 2023). Overlapping generations and the pioneer habit can thus slow down sex ratio selection in *Salix*. Indeed, simulation models, albeit for species with non-overlapping generations, suggest that populations with sex ratio distorter-restorer systems and female sex ratio bias might have an advantage in dispersal limited habitats, owing to their high seed production (Rood & Freedberg, 2016). Similarly, populations with female bias due to meiotic drive may have an advantage under strong competition (Unckless & Clark, 2014). We conclude that *Salix* species, including the arctic-alpine dwarf shrub *S. herbacea* investigated here, may have the ecological features that allow for the long-term persistence of dynamic and variable genetic sex ratio distortion systems that lead to female bias of the seed sex ratio.

## Conclusion

Our results suggest that *Salix herbacea* populations have a consistent female bias from the seed stage, likely caused primarily by genetic mechanisms. Population sex ratios are thus not much affected by changing environments, in contrast to other dioecious plant species where changing conditions have the potential to drive sex ratios to extremes and thereby endanger population persistence (Hultine et al., 2016; Petry et al., 2016). Variation in the secondary sex ratio biases (among flowering plants) between species has been attributed to sexual dimorphism in reproductive cost, for example, the high incidence of female bias in wind-pollinated species is interpreted as a result of higher reproductive costs in males leading to higher male mortality (Field et al., 2013). However, female-biased population sex ratios in seeds (primary sex ratio), and also variation in sex ratio between families, as reported here, are also consistent with entirely different mechanisms, such as meiotic drive and cyto-nuclear interactions (Rood & Freedberg, 2016; Unckless & Clark, 2014). These mechanisms arise from genetic conflict and can confer reproductive advantage in colonizing species (Rood & Freedberg, 2016; Unckless & Clark, 2014). Biases in the primary sex ratio are only rarely reported (Stehlik et al., 2008; Timerman & Barrett, 2019) but this may be because plant sex can only be inferred morphologically at flowering. We therefore conclude that biases in the primary sex ratio in dioecious plant species need to be considered more often in order to understand not only the underlying evolutionary and ecological mechanisms, but also potential risks to population persistence.

## Data availability statement

All data will be made available on Dryad upon acceptance.

## Author contributions

XM, AJC, CR and SK conceived the ideas and designed the methodology; AJC and CR collected the field data, XM and SK collected data from the controlled experiment; XM and SK analyzed the data; XM and SK led the writing of the manuscript. All authors contributed to the drafts and gave final approval for publication.

## Conflict of interest statement

All authors declare no conflict of interest.

## Acknowledgements

We are very thankful for the excellent work of Francesca Jaroszynska and Sven Buchmann in the field and of Nicolai Gmünder, Hanna Danko and Mattias Vass at our indoor experiment. We are much indebted to Mark van Kleunen and Julia Wheeler for allowing us to use their published and unpublished field data. This work was financed by a Sinergia grant of the Swiss National Science Foundation (CRSI33_130409/1) to CR, Mark van Kleuen and SK, by a project grant of the Swedish Research Council (Vetenskapsrådet, 2019-03844) to SK and by Uppsala University. Funding for open access publication was provided by Uppsala University.

## Notes

### Competing Interest Statement

The authors have declared no competing interest.

