## Supplementary material for "Female-biased population sex ratios caused by genetic rather than ecological mechanisms"

**Supplementary Table 1.** Numbers of seeds, seedlings, plants, and reproductive individuals in full-sib families of the dwarf shrub *Salix herbacea*, cultivated under controlled conditions. GH, greenhouse, GC, growth chamber

| Family | Seeds | Germinated seeds in Petri dish | Seedlings in randomized design, GH | Established seedlings in randomized design, GH | Plants transferred to GC, including replacements | Plants surviving to the end of the experiment | Females | Males | Flowering proportion | Sex ratio (proportion females) |
|---|---|---|---|---|---|---|---|---|---|---|
| **X02**[1] | 37 | 33 | NA | NA | 22 | 22 | 13 | 2 | 0.68 | 0.87 |
| **X05** | 97 | 93 | 91 | 62 | 64 | 63 | 7 | 21 | 0.44 | 0.25 |
| **X06** | 59 | 57 | 53 | 24 | 24 | 23 | 9 | 7 | 0.7 | 0.56 |
| **X07**[2] | 2 | 2 | 2 | 1 | 1 | 1 | 0 | 1 | 1 | 0 |
| **X08** | 148 | 127 | 84 | 54 | 69 | 63 | 13 | 36 | 0.78 | 0.27 |
| **X09** | 148 | 146 | 138 | 65 | 65 | 64 | 38 | 3 | 0.64 | 0.93 |
| **X10** | 80 | 67 | 57 | 21 | 21 | 21 | 9 | 5 | 0.67 | 0.64 |
| **X11**[3] | 132 | 130 | 116 | 78 | 82 | 79 | 56 | 11 | 0.85 | 0.84 |
| **X12** | 117 | 111 | 111 | 51 | 51 | 51 | 18 | 20 | 0.75 | 0.47 |
| **X13** | 15 | 13 | 13 | 8 | 8 | 8 | 7 | 0 | 0.88 | 1 |
| **X14** | 173 | 164 | 155 | 48 | 50 | 50 | 21 | 16 | 0.74 | 0.57 |
| **X15**[2] | 5 | 4 | 4 | 3 | 3 | 3 | 0 | 0 | 0 | 0 |
| **X16** | 12 | 11 | 10 | 6 | 6 | 6 | 3 | 2 | 0.83 | 0.6 |

| | | | | | | | | | | |
|---|---|---|---|---|---|---|---|---|---|---|
| **X17** | 25 | 24 | 22 | 13 | 13 | 12 | 11 | 0 | 0.92 | 1 |
| **X18**[2] | 4 | 3 | 3 | 1 | 1 | 0 | 0 | 0 | NA | 0 |
| **X19** | 35 | 34 | 33 | 19 | 20 | 19 | 14 | 1 | 0.79 | 0.93 |
| **X20** | 54 | 52 | 50 | 32 | 32 | 30 | 22 | 0 | 0.73 | 1 |
| **X22** | 49 | 46 | 44 | 27 | 29 | 23 | 20 | 0 | 0.87 | 1 |
| **X23** | 28 | 27 | 27 | 8 | 8 | 8 | 4 | 1 | 0.63 | 0.8 |
| **X25** | 38 | 37 | 36 | 17 | 18 | 18 | 14 | 3 | 0.94 | 0.82 |
| **X27** | 19 | 16 | 15 | 13 | 14 | 13 | 9 | 2 | 0.85 | 0.82 |
| **X29** | 13 | 12 | 12 | 8 | 8 | 8 | 4 | 2 | 0.75 | 0.67 |
| **X30** | 6 | 5 | 4 | 3 | 4 | 4 | 3 | 1 | 1 | 0.75 |
| **X31**[4] | 10 | 10 | 10 | 8 | 8 | 7 | 8 | 0 | 1 | 1 |
| **X32** | 65 | 64 | 62 | 44 | 44 | 44 | 39 | 0 | 0.89 | 1 |
| **X34** | 25 | 22 | 21 | 10 | 11 | 11 | 8 | 1 | 0.82 | 0.89 |
| **X36** | 60 | 58 | 57 | 36 | 37 | 34 | 20 | 11 | 0.91 | 0.65 |
| **X39**[5] | 35 | 31 | 29 | 19 | 21 | 20 | 12 | 5 | 0.85 | 0.71 |
| **X40** | 30 | 29 | 29 | 23 | 23 | 23 | 17 | 1 | 0.78 | 0.94 |

[1] Family X02 was planted as a trial and placed on an extra table.
[2] Three families (X07, X15, X18) with fewer than 3 surviving plants were removed from the regression analysis.
[3] One female of X11 was sampled in growth period 3 (unrelated experiment).
[4] One female of X31 died in growth period 3.
[5] One individual of X39 had deformed sexual organs that combined male and female traits and was excluded from further analyses.

**Supplementary Table 2.** The number of reproductive and non-reproductive individuals of the dwarf willow *Salix herbacea* in a controlled experiment over three growth periods and overall.

| Growth period | Females | Males | Vegetative | Flowering proportion | Sex ratio (proportion females) |
|---|---|---|---|---|---|
| 2 | 198 | 81 | 466 | 0.37 | 0.71 |
| 3 | 228 | 97 | 411 | 0.44 | 0.7 |
| 4 | 354 | 137 | 237 | 0.67 | 0.72 |
| All | 398 | 152 | 178 | 0.76 | 0.72 |

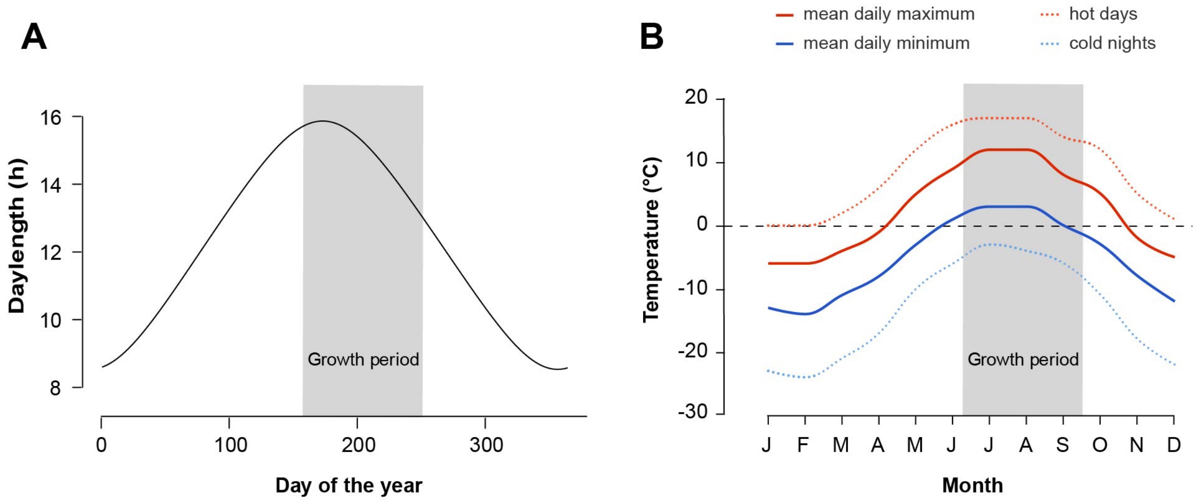


**C** **Experimental conditions**

| | | Spring-summer | Autumn | Winter |
|---|---|---|---|---|
| **Duration (days)** | Growth period 1 | 49-81 | 70-81 | 101 |
| | Growth period 2 | 35 | 71 | 92 |
| | Growth period 3 | 34 | 73 | 111 |
| | Growth period 4 | 39 | 70 | 275 |
| **Temperature** | | day: 8-16℃<br>night: 4-10℃ | day: 16-6℃<br>night: 10-4℃ | 0℃ |
| **Day length** | | 14-16h | 16-12h | 0 |
| **Light intensity (PAR)** | | 150 $\mu Em^{-2}s^{-1}$ | 150-100 $\mu Em^{-2}s^{-1}$ | 0 |

**Supplementary Figure 1.** Climatic conditions in a natural population of the dwarf willow *Salix herbacea* at Jakobshorn Switzerland (**A**, **B**) and in an experiment under controlled conditions (**C**). **A**, daylength obtained through the R package *geosphere* (Hijmans, R., 2022, geosphere: Spherical Trigonometry. R package version 1.5-18), **B**, mean daily minima and maxima of air temperature overall and for hot days and cold nights (1990-2020); data from *meteoblue.com* (accessed 15. November 2023) and used with permission. **C**, duration, temperature, day length and light intensity in simulated seasons.
